# *PhageDPO*: Phage Depolymerase Finder

**DOI:** 10.1101/2023.02.24.529883

**Authors:** Maria Vieira, José Duarte, Rita Domingues, Hugo Oliveira, Oscar Dias

## Abstract

Bacteriophages are the most predominant and genetically diverse biological entities on Earth. They are bacterial viruses which encode numerous proteins with potential antibacterial activity. However, most bacteriophage-encoded proteins have no assigned function, hindering the discovery of novel antibacterial agents. In particular, there has been a growing interest in exploring recombinant bacteriophage depolymerases from the fundamental standpoint, but mostly for biotechnological applications to control bacterial pathogens. Due to the lack of efficient identification tools, we developed *PhageDPO*, the first developed tool that predicts depolymerases in bacteriophage genomes using machine learning methods.

**Availability and implementation:** *PhageDPO* was integrated into a Galaxy framework available online at: bit.ly/phagedpo.

## 1. Introduction

Bacteriophages (phages) are viruses that infect and replicate within bacteria (Duckworth and Gulig 2002). Generally, phages recognize bacterial hosts through receptor-binding proteins (RBPs). In several phages, these RBPs encode enzymes, which facilitate viral binding and degradation of bacterial carbohydrates (e.g., capsules, lipopolysaccharides), called depolymerases (DPOs). Recombinant DPOs have been studied, by leveraging this function, to remove bacterial carbohydrates and turn bacterial pathogens less virulent, thus more easily controlled by the host immune system (Oliveira, Costa, et al. 2019; Oliveira, Mendes, et al. 2019). A recent review on the diverse biotechnological applications of phage DPOs can be accessed here (Oliveira, Drulis-Kawa, and Azeredo 2022).

Given that phages are the most abundant biological in the biosphere, with an estimated 10^31^ phages and outnumbering bacteria by ten-fold, they encode an endless arsenal of proteins, such as DPOs, which might be used for biotechnological applications. Nevertheless, efficient annotation tools are needed to ease the identification of DPOs, which are amongst the most diverse proteins in the phage proteome. Current DPO identification is limited to manual and homology-based tedious processes. Latka *et al*. (Latka et al. 2019) described the identification of DPOs in specific *Klebsiella* phage genomes, by filtering phage RBPs and then applying consecutive homology-based rules spanning BlastP (Altschul et al. 1990), Phyre2 (Kelley et al. 2015), SWISS-MODEL (Bordoli and Schwede 2011), HMMER (Finn, Clements, and Eddy 2011), and HHPred (Soding, Biegert, and Lupas 2005). Based on these tools, the authors selected a range of criteria that a protein must have to present a putative DPO activity. These criteria included: size (>200 residues), annotation (tail fibre/tail fiber/tail spike or hypothetical protein in the NCBI database), and homologies to known enzymatic domains (lyase or hydrolase). The length of homology with one of these enzymatic domains should span at least 100 residues and a typical β-helical structure should be predicted by Phyre2. With this approach, several putative DPOs were predicted. However, such efforts were the result of extensive manual curation and the predicted DPOs were not experimentally validated. Therefore, such an approach does not provide a user-friendly tool capable of predicting phage DPOs.

Due to the lack of bioinformatics tools to identify these proteins, *PhageDPO* was developed. This tool, based on machine learning methods, explores the whole genome to find phage DPOs and returns the percentage of positive predictions for phage DPO.

## 2. Materials and Methods

### Data

*PhageDPO* was trained with phage DPOs retrieved from the National Center for Biotechnology Information (NCBI)’s Protein database (Supplementary Table S1 and Table S2). The DPOs were collected based on a filtered search by proteins including at least one of six DPO-associated domains (cl40625, cd20481, Pfam12219, cl22684, Pfam12217, Pfam13472) or a constrained query performed through NCBI’s Entrez Programming Utilities, described in Supplementary Table S3.

This process returned 1437 sequences for positive cases and 22976 sequences for negative cases. To test the influence of negative cases on model performance, two datasets were created with a different number of negative cases, one with 2874 cases and the other with 5748 cases, indicative of dataset d4311 (1437 positive + 2874 negative) and d7185 (1437 positive + 5748 negative), respectively.

### Features

Based on sequence properties, 578 features were calculated. Data features included physicochemical characteristics, such as length, aromaticity, isoelectric point, secondary structure fraction, Composition Transition Distribution (CTD) and features based on amino acid composition. The full set of features is available in the supplementary data.

### Models

Two algorithms, Support Vector Machines (SVMs) and Artificial Neural Networks (ANNs), were used to train machine learning models to predict phage DPOs. The hyperparameters tested in each model are described in Supplementary Table S4.

The models selected for integration in the tool were the SVM model trained with dataset d4311 and the ANN model trained with dataset d7185.

### Experimental Validation

*PhageDPO* was assessed against two datasets: i) phage genomes with known and validated DPOs and ii) phage genomes without known DPOs (novel phages). In the latter dataset, the DPOs were predicted and subsequently validated *in vitro*.

## 1. Application

### Results

The SVM model exhibited an accuracy of 95%, a precision of 98% and a 91% recall, whereas the ANN model showed an accuracy of 98%, 99% of precision and 96% of recall. The SVM model seems to perform better in predicting true DPO sequences and preventing false positives, while the ANN model on ensuring that all DPO sequences are identified. These and other results are detailed in Supplementary Table S5 and S6.

### Galaxy Implementation

*PhageDPO* was developed in Python 3.7 and implemented in the Galaxy framework, providing a user-friendly graphical interface. The tool can be found in the Phage Annotation side-left bar and requires as input a FASTA file format with the nucleotide sequences of the ORFs. As an advanced option, users can select the model to run, considering that SVM (by default) will return fewer predictions than the ANN model, but with a high probability of being actual DPOs. *PhageDPO* returns an HTML table with sequence identification and the respective score of positive prediction. Case studies conducted on i) phage genomes with validated DPOs in literature and ii) novel phages, followed by experimental validation of DPO activity performed in this study, demonstrated that *PhageDPO* has a good performance in predicting DPO sequences (Supplementary Table S7 and S8).

## 2. Conclusion

*PhageDPO* is the first software tool that uses machine learning to predict phage DPOs. Despite having tested several models during the development of *PhageDPO*, the ANN and SVM achieved the best results, with small differences, as the SVM model returns fewer predictions, but with a high probability of being DPOs. Moreover, the tool performed well when its predictions were assessed in laboratory experiments. Generally, this tool provides good model performance, making the task of finding a DPO in phage genomes, easier, faster and more accurate.

## Acknowledgements

We thank Gregory Resch for sharing the *Acinetobacter* phages used in this study.

