## Supplementary Table for "*PhageDPO*: Phage Depolymerase Finder"

### 1. Data

All the data obtained for this work was gathered in August 2021. Six depolymerases (DPO)-associated domains (Table S3) were used during the construction of positive dataset within National Center for Biotechnology (NCBI)'s Conserved Domains Database (CDD) and their related proteins. Accordingly, 1751 DPO sequences were obtained.

*Table S3. Domains associated with DPOs activity*

| Domain | Sequences obtained | Associated domain and protein (example) |
|---|---|---|
| **cl40625** | 517 | YP_007349010 |
| **cd20481** | 287 | ASN73504 |
| **Pfam12219** | 111 | YP_338127 |
| **cl22684** | 68 | CBY99579 |
| **Pfam12217** | 75 | YP_338127 |
| **Pfam13472** | 693 | ARB10970 |

NCBI's protein database was used to ensure that all DPOs were collected, even those not present in CDD. The constraints for the query created to obtain positive cases were the following:

*viruses[porgn: txid28883]*
*AND tail**
*AND (*glycanase*[Text Word]*
*OR *alginate*[Text Word]*
*OR *rhamnosidase*[Text Word]*
*OR *hyaluronidase*[Text Word]*
*OR *hyaluronate*[Text Word]*
*OR *eps-degrading*[Text Word]*
*OR *levanase*[Text Word]*
*OR *dextranase*[Text Word]*
*OR *xylanase*[Text Word])*

This query allowed retrieving 677 additional sequences for a total of 2428 positive sequences. CD-HIT (Fu et al., 2012) was used to reduce redundant data, resulting in 1437 positive DPO sequences.

Likewise, negative cases were obtained using the following query constraints:

*(viruses[porgn: txid28883]*
*AND srcdb refseq[Properties])*
*NOT lyase*[Text Word]*
*NOT pectate*[Text Word]*
*NOT pectin*[Text Word]*
*NOT depolymerase*[Text Word]*
*NOT glycanase*[Text Word]*
*NOT endoglycosidase*[Text Word]*
*NOT alginate*[Text Word]*
*NOT rhamnosidase*[Text Word]*

*NOT hyaluronidase*[Text Word]*
*NOT hyaluronate*[Text Word]*
*NOT eps-degrading*[Text Word]*
*NOT levanase*[Text Word]*
*NOT dextranase*[Text Word]*
*NOT xylanase*[Text Word]*
*NOT pfam12708*[Text Word]*
*NOT cd20481*[Text Word]*
*NOT pfam12219*[Text Word]*
*NOT cl22684*[Text Word]*
*NOT sgnh hydrolase*[Text Word]*
*NOT gdsl hydrolase*[Text Word]*
*NOT pfam12217*[Text Word]*
*NOT pfam13472*[Text Word]*

For negative cases, the results were limited to the first 30000 entries. The results were curated through keyword check, and duplicate sequences removed using CD-HIT (Fu et al., 2012), resulting in 22976 negative DPO sequences.

#### 2. Feature Calculation

Features related to the protein sequences' properties were calculated for model training purposes. These features can be divided into:

- **Amino acid composition** (**AAC**) and **Nucleotide composition** (**NC**): **AAC** and **NC** are composed of 20 and 4 vectors, respectively. The formulation of these compositions is indicated below in Equation 1.

$$Comp_j = \sum_{i=1}^{l} \sigma_i \ 1 \tag{1}$$

  Where $Comp_j$ represents the composition of $j$ (amino acid or nucleotide) and $l$ the sequence length. The vector is constructed based on two conditions: $\sigma_i = 1$, if the $i$ occurrence is j-type, and $\sigma_i = 0$, if the $i$ occurrence is not j-type. The variable $j$ range from 1 to 20 in protein sequences and 1 to 4 in DNA sequences.

- **Aromaticity**, **Secondary Structure fraction,** and **Isoelectric point**: The calculation of **Aromaticity** and **Isoelectric point** results in one vector each. For the **Secondary Structure** fraction, 3 vectors are calculated.

  The calculation of **Aromaticity** returns the relative frequency of aromatic amino acids. The formula for this calculation is shown below in Equation 2.

$$Aromaticity = \sum_{i=1}^{20} \gamma_i f_i \tag{2}$$

Where $f_i$ represents the frequency of amino acid $i$. $\gamma_i = 1$, if the amino acid is Phe, Tyr, or Trp, otherwise $\gamma_i = 0$.
The **Isoelectric point** represents a vector of pH where the net charge of the protein is zero.
The **Secondary Structure** fraction is based on the calculation of the fraction of amino acids that tend to be represented as one of the secondary structures: $\alpha$-helix, $\beta$-sheet, and *turns*. For each structure, Equations 3, 4, and 5 respectively, can be seen below.

$$Helix = \sum_{i=1}^{20} \alpha_i f_i \quad (3)$$

$$Turns = \sum_{i=1}^{20} \theta_i f_i \quad (4)$$

$$Sheets = \sum_{i=1}^{20} \beta_i f_i \quad (5)$$

Where $f_i$ represents the frequency of amino acid $i$ type. Each vector is constructed based on these conditions:
$\alpha_i = 1$, if the amino acid is Val, Ile, Tyr, Phe, Trp, or Leu, otherwise $\alpha_i = 0$
$\theta_i = 1$, if the amino acid is Asn, Pro, Gly, or Ser, otherwise $\theta_i = 0$
$\beta_i = 1$, if the amino acid is Glu, Met, Ala, or Leu, otherwise $\beta_i = 0$

- **Composition Transition Distribution** (CTD): The calculation of CTD results in a vector of 147 features. This metric clusters amino acids into three classes. Properties of amino acids, such as polarity, hydrophobicity, charge, solvent accessibility, secondary structure, and normalized van der Waals volume, were used. The metric can be divided into three categories: **Composition** - which represents the composition of amino acid in each property divided by the total number of amino acids. **Transition** - measures how often amino acids of one property are followed by amino acids of another property. **Distribution** - measures length, where the first 25, 50, 75, and 100% of the amino acids from a particular property are located, respectively.

The **Composition** for each property is given according to Equation 6.

$$Composition_j = \frac{1}{l} \sum_{i=1}^{l} \sigma_i \quad (6)$$

Where $Comp_j$ represents the composition of amino acids for each property $j$=1, 2, 3 for the sequence length $l$. This vector, like the amino acid composition calculation, is built based on a condition: $\sigma_i = 1$, if the $i$ occurrence is j-type, and $\sigma_i = 0$, if the $i$ occurrence is not j-type.

The **Transition** for each property is calculated according to Equation 7.

$$Transition_{mn} = \frac{D_{mn} + D_{nm}}{l - 1} \tag{7}$$

Where $mn$ "12" "13", "23" and $D_{mn}$ and $D_{nm}$ are the numbers of dipeptides encoded as "$mn$" and "$nm$" in the sequence, respectively.

Equation 8 describes the **Distribution** of properties along the protein chain.

$$R_{j,p} = \frac{P}{100}\sum_{i=1}^{l} \sigma_i \tag{8}$$

Where $R_{j,p}$ represents the number of residues for group $j$ within a given percentage $P$ for a protein length $l$, where $P$ = 1, 25, 50, 100 and $j$ = 1,2,3.

- **Dipeptide Composition** (**DPC**). This calculation generates 400 features for each protein sequence. **DPC** is calculated according to Equation 9 below.

$$DPC_j = \frac{D_j}{l - 1} \tag{9}$$

Where the variable $j$ ranges values from 1 to 400.

### 3. Feature Selection

Recursive Feature Elimination (RFE) was used as a method for feature selection. After applying this method, some features are eliminated based on weights provided by the algorithm.

### 4. Model Optimization

Support Vector Machines (SVM) and Artificial Neural Networks (ANN) were the selected machine learning algorithms.

For both algorithms, the models were trained with datasets containing 1437 positive cases. However, regarding the negative case, datasets of different sizes were tested. Hence, the first dataset encompassed 2874 and the second dataset accounted for 5748 negative cases, corresponding to dataset d4311 and d7185, respectively. The models trained with these datasets were optimized through a Grid (hyperparameter) Search. The training details are available in Table S4.

Table S4. Hyperparameters selected for each model in the Grid Search optimization

| Model | Hyperparameters | |
|---|---|---|
| ANN | d4311 | Activation function: Hyperbolic Tangent (tanh)<br>Solver for weight optimization: Adam<br>alpha: 0.001<br>One hidden layer with 50 neurons |
| | d7185 | Activation function: Rectified Linear Unit Layer (ReLU)<br>Solver for weight optimization: Adam<br>alpha: 0.0001<br>One hidden layer with 100 neurons |
| SVM | d4311 | C (regularization): 10<br>gamma: 0.1<br>kernel: radial basis function (rbf) |
| | d7185 | C (regularization): 10<br>gamma: 0.01<br>kernel: radial basis function (rbf) |

### 5. **Benchmarking**

Model performance was evaluated using 5-fold cross-validation. The assessed metrics were Percentage of Examples Correctly Classified (PECC), precision, and recall. For each model and dataset, a confusion matrix was created. The results are shown in Tables S5 and S6.

Table S5. Confusion matrices of the ANN models for both datasets

| Real \ Predicted | Positive | Negative | Total |
|---|---|---|---|
| Positive | 1305 | 132 | 1437 |
| Negative | 121 | 2753 | 2874 |
| Total | 1426 | 2885 | 4311 |

| Real \ Predicted | Positive | Negative | Total |
|---|---|---|---|
| Positive | 1281 | 156 | 1437 |
| Negative | 110 | 5638 | 5748 |
| Total | 1391 | 5794 | 7185 |

Table S6. Confusion matrices of the SVM models for both datasets

| Real \ Predicted | Positive | Negative | Total |
|---|---|---|---|
| Positive | 1273 | 164 | 1437 |
| Negative | 58 | 2816 | 2874 |
| Total | 1331 | 2980 | 4311 |

| Real \ Predicted | Positive | Negative | Total |
|---|---|---|---|
| Positive | 1282 | 155 | 1437 |
| Negative | 89 | 5659 | 5748 |
| Total | 1371 | 5814 | 7185 |

#### 5.1 Case Studies

For phage genomes with known DPOs, the predictive models were applied to three phage genomes: 1) *Acinetobacter* phage vB_Api_3043-K38 (accession number: MZ593174); 2) *Klebsiella* phage RAD2 (accession number: NC055956); 3) and 4) *Escherichia* phage vB_EcoP_G7C (accession number: NC015933). The respective DPOs encoded by each phage were experimentally validated before (Domingues et al., 2021; Dunstan et al., 2021; Prokhorov et al., 2017). The predicted *PhageDPO* results on these genomes are summarized in Table S7.

*Table S7. DPO prediction for both SVM and ANN models. * Only significant results above 70 % are shown.*

| Phage | Model | Protein Identifier | DPO Prediction (%)* |
|---|---|---|---|
| *Acinetobacter* phage 3043-K38 | SVM | QYC50642 | 99.0 |
| | ANN | QYC50642 | 100.0 |
| *Klebsiella* phage *RAD2* | SVM | YP_010115729 | 99.0 |
| | | YP_010115728 | 81.0 |
| | ANN | YP_010115729 | 100.0 |
| | | YP_010115728 | 78.0 |
| *Escherichia* phage G7C | SVM | YP_004782196 | 100.0 |
| | | YP_004782195 | 94.0 |
| | ANN | YP_004782196 | 100.0 |
| | | YP 004782195 | 92.0 |
| | | YP 004782143 | 90.0 |

The results obtained by *PhageDPO* were in accordance to the literature. *Acinetobacter* phage 3043-K38 encodes a single DPO that degrades capsules of *Acinetobacter baumannii* (QYC50642), as predicted with high probability for both SVM and ANN models (Domingues et al., 2021). *Klebsiella* phage RAD2 also encodes a single DPO (YP 010115729) that targets the capsule of the *Klebsiella pneumoniae* (Dunstan et al., 2021). The SVM model predicted the correct protein with 99% probability, while the ANN model predicted the same protein with 100% probability. Both models also predicted, with lower percentage, a false positive protein (YP 010115728), annotated as a putative tail spike. The *Escherichia phage* G7C has a new kind of DPO (YP 004782195), that modifies instead of degrading the lipopolysaccharide (Prokhorov et al., 2017). The SVM model predicted the correct protein (YP 004782195) with 94% probability and a second one (YP 004782196) with 100% probability. The ANN model predicted the same proteins (YP 004782195.1, YP 004782196) with 100% and 92% respectively, and third (YP 004782143) with 90% probability. The protein YP 004782195, predicted by both models, is annotated as a tail fiber. Further lab tests should be performed to find if this phage encodes two DPOs, as suggested by the predictive model. Protein YP 004782143, predicted by the ANN model, is annotated as RNA polymerase, and therefore, a false positive.

The predictive models were likewise applied to six novel phage genomes without known DPO, of different taxonomies i.e. phylogenetic distinct:

- *Acinetobacter* phage vB_Ab_3042-K38 (*Obolenskvirus* genus)
- *Acinetobacter* phage vB_Ab_3073b-K32 (*Obolenskvirus* genus)
- *Acinetobacter* phage_vB_Ab_3073a-K32 (unclassified genus)
- *Acinetobacter* phage vB_AbP_F70-K44 (*Friunavirus* genus)

- *Acinetobacter* phage_vB_AbP_P2 (*Friunavirus* genus).
- *Acinetobacter prophage* (unclassified genus)

The results of *PhageDPO* performed in all genomes are summarized in Table S8.

*Table S8. DPO prediction for both SVM and ANN models. * Only significant results above 70 % are shown.*

| Phage | Model | Protein Identifier | DPO Prediction (%) * |
|---|---|---|---|
| *Acinetobacter phage* 3042-K38 | SVM | OQ427159 | 100.0 |
| | ANN | | 100.0 |
| *Acinetobacter phage* 3073b-K32 | SVM | OQ427162 | 100.0 |
| | ANN | | 100.0 |
| *Acinetobacter phage* 3073a-K32 | SVM | OQ427161 | 100.0 |
| | ANN | | 100.0 |
| *Acinetobacter phage F70-K44* | SVM | OQ378314 | 99.0 |
| | ANN | | 100.0 |
| *Acinetobacter phage P2* | SVM | ASN73558 | 100.0 |
| | ANN | | 100.0 |
| *Acinetobacter prophage* | SVM | WP_000729646 | 99.0 |
| | ANN | | 97.0 |

Predicted DPOs encoding genes were cloned into *E. coli* pET28a plasmids and the resulting proteins overexpressed and purified using affinity chromatography, as previously described (Oliveira et al., 2017). Finally, drops of purified proteins were spotted on bacterial lawns that confirmed the depolymerase activities by visual confirmation of haloes (Figure S1). All, except the DPO from the prophage were confirmed experimentally. The fact that a high prediction score was obtained for the prophage protein, but no depolymerase activity was observed, can indicate that i) it could be a false result, or ii) the DPO is active in another bacterial host other that the one tested, as DPOs are known to be very specific proteins.

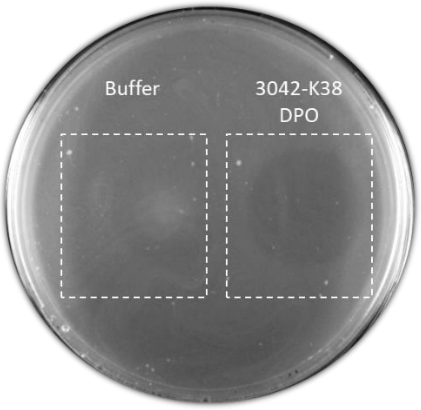

*Figure S1. Confirmation of depolymerase activity predicted in PhageDPO.* This is an example of visual inspection of DPO activity performed using the predicted *Acinetobacter* phage 3042-K38 DPO.
